# Interplay of mechanics, growth and viral infection dynamics in a spatially expanding bacterial colony

**DOI:** 10.64898/2026.10.02.755697

**Authors:** Godwin Stephenson, Amit Shimoga Nadig, Toshi Parmar, K Vijay Kumar, Namiko Mitarai, Sandeep Krishna, Shashi Thutupalli

**Author notes:** These authors contributed equally to this work (Godwin Stephenson, Amit Shimoga Nadig).

## Abstract

We investigate the dynamics of the spread of the infection due to a phage-*λ* during the range expansion of an *E coli* colony. We find a rich panoply of spatio-temporal dynamics at large colony-level length scales, arising due to the interplay of mechanical forces, growth and viral infection dynamics at smaller cellular length scales. Strikingly, the phages “surf” the front of the growing bacterial colony by hitchhiking on *E coli* that are advected due to the growth of bacteria in the colony, resulting in an anisotropic spread of the phages. We identify microscopic processes – of the phage release during lysis (occurring on short millisecond timescales at length scales close to that of a single bacterium) and local nematic lignment of the rod-like *E coli* bacteria (occurring on timescales comparable to the bacterial growth rate and the length scales of a few bacterial cells) – that enhance the advective effects driving the hitchhiking behavior. Combining our experiments with a mathematical model, we explain the multiple spatio-temporal dynamical regimes – from the coexistence of uninfected bacteria, resistant (lysogenic) cells and phages, to fixed points where the entire population turns resistant or remains uninfected. Our work broaches a new frontier in the investigation of eco-evolutionary processes arising from the mechanical and ecological forces due to the interactions between viruses and their hosts.

## 1 Introduction

Growth and dispersal, accompanied by spatial range expansion, are a fundamental occurrence in the eco-evolutionary history of many species, from microbes to human populations [1]. In natural settings, such range expansions invariably involve inter-species interactions, be it the competition between invading and native species [2], the spread of an infectious species (pathogen) within another (host) or even the mutualistic colonisation of spatial niches [3]. Measuring and predicting the outcome of such spatio-temporal dynamics in natural environments has remained difficult — the spread of infections, for example, poses a continually evolving challenge due to dynamic changes in the host population structure and mechanisms of pathogen spread [4]; laboratory studies using microbial model systems, on the other hand, have yielded many insights [5, 6, 7]. Specifically, the growth driven expansion of microbial populations has been used to uncover a spatial demixing (sectoring) of “genotypes”, reflecting an underlying competition between the effects of genetic drift and selection [5, 8, 9]. The resultant spatial structure has strong consequences for population outcomes such as the frequencies of mutant clones [10], resistance to drugs [11] and maintenance of co-operation [12]. Apart from ecological forces such as interspecies competition, fluctuations, both intrinsic and extrinsic [13], and mechanical interactions [14] shape the eco-evolutionary dynamics. Here, we exploit one of the best studied inter-species microbial interaction — that between viruses and bacteria — to study the population dynamics driven by the interplay between infection dynamics, growth driven range expansion and the associated mechanics.

Virus-bacteria interactions predominantly occur in spatial and structured natural environments ranging from the human gut to soils and oceans [15, 16, 17]. However, despite the pervasiveness of such spatial settings, most of the investigations of their eco-evolutionary dynamics are either in well-mixed conditions [18] or in the relatively simple spatial setting of a so-called ‘plaque assay’ *i*.*e*. the spread of phage infection in a uniform background of bacteria [19, 20]. In such a plaque assay, both virulent phage — that kill bacteria via “lysis” — and temperate phage — that exhibit “lysis” and “lysogeny” — spread in an isotropic, wave-like manner resulting in circular bacterial clearings *i*.*e*. plaques. While lysis results in bacterial death via a bursting of the cell and release of freshly replicated virions, lysogeny refers to a dormant state that temperate phage can enter when they infect bacteria: the viral genetic material integrates within the genome of the bacteria or forms a replion and does not replicate as phage; the bacteria however replicate, together with the viral genetic material, and are resistant to further infection of the same virus. Thus, while virulent phage produce ‘clear plaques’, temperate phage leave behind some lysogenic bacteria in the cleared area resulting in ‘turbid plaques’; these dynamics have been extensively studied [21, 22, 23, 24, 25, 26]. In more complex spatial settings, one may imagine that the spread of virulent phage resembles that of any other infectious agent amongst a host population [27, 28]. However, the spread of temperate phage, which involves the two infection outcomes, *i*.*e*., lysis and lysogeny, may lead to more complex spatio-temporal demographic patterning. The choice between these states depends on multiple intrinsic and extrinsic factors (*e*.*g*. environmental, bacterial densities, growth rates etc.). For *E. coli* infected by phage *λ*, the most studied temperate phage, one of the more important factors is the Average Phage Input (API) which is the ratio of number of viruses to the number of bacteria in the medium. As shown by Zeng, Lanying et.al. the lysogenization frequency is a function of API and increases with increase in number of phages infecting the bacterium. [29, 30, 31, 32, 33, 34]. Temperate viral dynamics in bacterial populations therefore includes uninfected and resistant (lysogenic) cells in addition to those that die via lysis and produce fresh viruses — this well established overall demographic structure therefore provides a fertile setting in which to investigate their growth-driven spatio-temporal dynamics.

We perform experiments using *E coli* bacterial colonies that grow spatially outward, driven by a self-organized nutrient gradient, with phage-*λ* mixed into this expanding colony of bacteria. We find that the dynamic interplay between single cell-level growth and mechanics couple with phage transport and infection processes at larger length-scales causing the phages to “surf” the propagating bacterial front. The surfing phages drive an anisotropic spread of the infection, along the direction of the colony growth, thereby forming elongated “infection rivers”; the local mechanics of bacterial growth and interactions cause the cells to orient themselves along these rivers, furthering their propagation. The meandering and spatial organization of these rivers are not only governed by these mechanics but also the differential transport of phages within and outside the bacterial colonies and by the initial conditions. Altogether, this leads to a rich tapestry of spatio-temporal demographic patterning. We summarize the findings into a mathematical model that takes into account the cell-growth mediated propagation (advection) of infected cells and phages and demonstrate that it captures the basic features of the observed patterns. Our results not only have consequences for understanding phage–bacterial dynamics in natural settings but also for emergent population structures and organization due to the spatial co-expansion of interacting species [5, 10].

## 2 Results

### 2.1 Co-existence and extinction in viral infections in spatially growing bacterial colonies

The experimental system that we use [29] allows us to visualise the phages and also the different infection states of the *E coli* using fluorescence microscopy (Figure 1**A**): briefly, the phage capsid decorative protein, gpD is translationally fused with the fluorescent mTurquoise2 (blue) molecule so that single free phages appear as bright blue spots. Upon lytic decisions, the cells produce progeny phages and the cells appear full of blue spots in the lytic state. Lysogenic reporters are constructed by transcriptionally fusing mKate2 (Red) to the phage lytic repressor gene, cI. Upon lysogenic decisions, cells express the red lysogenic reporter as they grow and divide. We seed an initial circular bacterial colony (~ 10^5^ cells in exponential phase) along with a small proportion of phages (in this case ~ 10 phages). The outcome of the infections that follow is a dynamic interplay of viral and bacterial transport and growth, generating distinct patterns of infection which are closely coupled to the bacterial background (Figure 1**B**). These patterns are different from the more commonly studied isotropic, roughly circular turbid plaques formed when temperate phage are spotted on a uniform bacterial lawn (Figure SI 5**A**).

**Figure 1:**
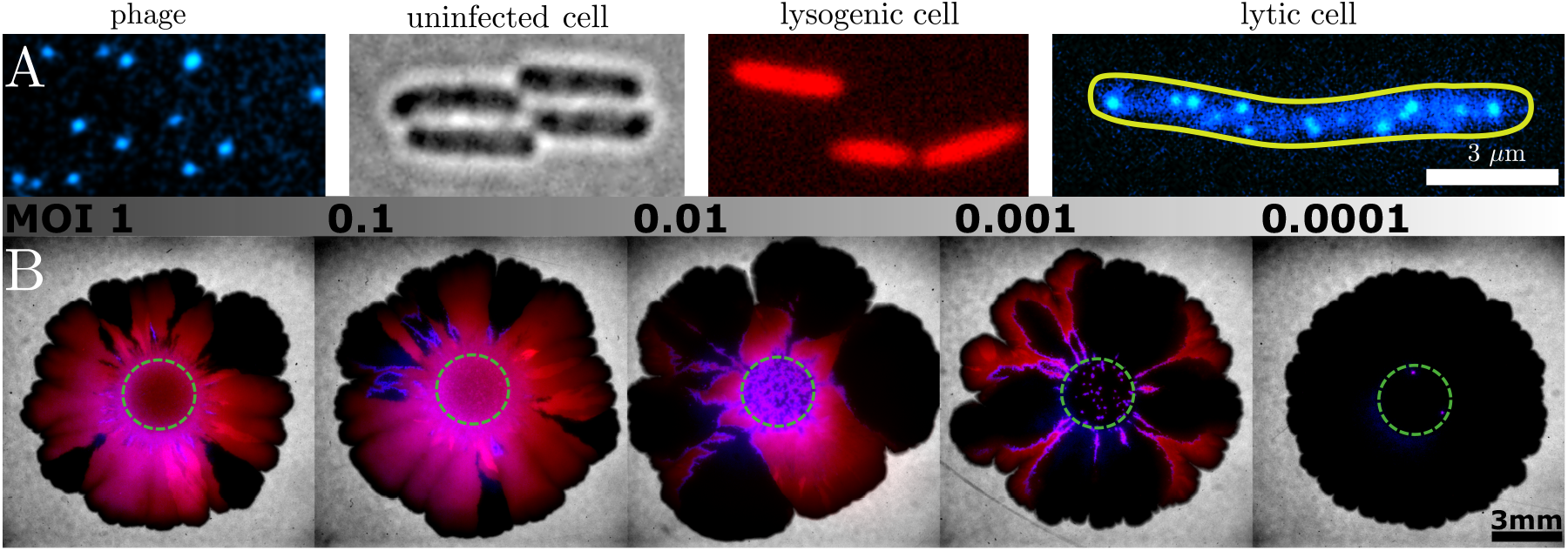
**A**. Phages with gpD capsid proteins fused with mTurquoise2 enable individual visualization as bright fluorescent spots. Brightfield image of unlabeled uninfected cells. Multi-infected cells expressing mKate2 on entering lysogeny. Elongated single-infected lytic cells, revealing the internal assembly of phages. **B**. Colony infection patterns. Fluorescence and bright field overlay of the infected cell colonies from 1 to 10^*−*4^ API with dashed circles indicating the homeland (Scale bar = 3 mm). At high APIs, the colony is dominated by lysogenic bacteria owing to a large phage output during the second wave of infection. While at low APIs the number of phages infecting is not sufficient to outgrow bacterial growth and hence uninfected cells prevails. At an intermediate range of APIs the uninfected bacteria and lysogenic bacteria co-exist with the phage infection surfing the colony front.

### 2.2 Local ordering of cells near the growth front and in lytic rivers

*E. coli* are rod-shaped bacteria (here average aspect ratio of 4:1) that elongate along their longitudinal cell axis before dividing mid-axially into two daughter cells. At the edge of a colony (Figure 2**A**), this growth induces the alignment of neighbouring cells, orienting themselves perpendicular to the direction of growth of the colony front [7]. This orientation persists in a region of thickness ~2-3 cell widths. Beyond that there is a region, of thickness ~2-3 cell *engths*, where cells are oriented in a direction parallel to the growth, i.e., their long-axis points approximately radially outward from the homeland. Further in, the orientation of cells is more disordered, with small domains exhibiting local order in effectively random directions.

**Figure 2:**
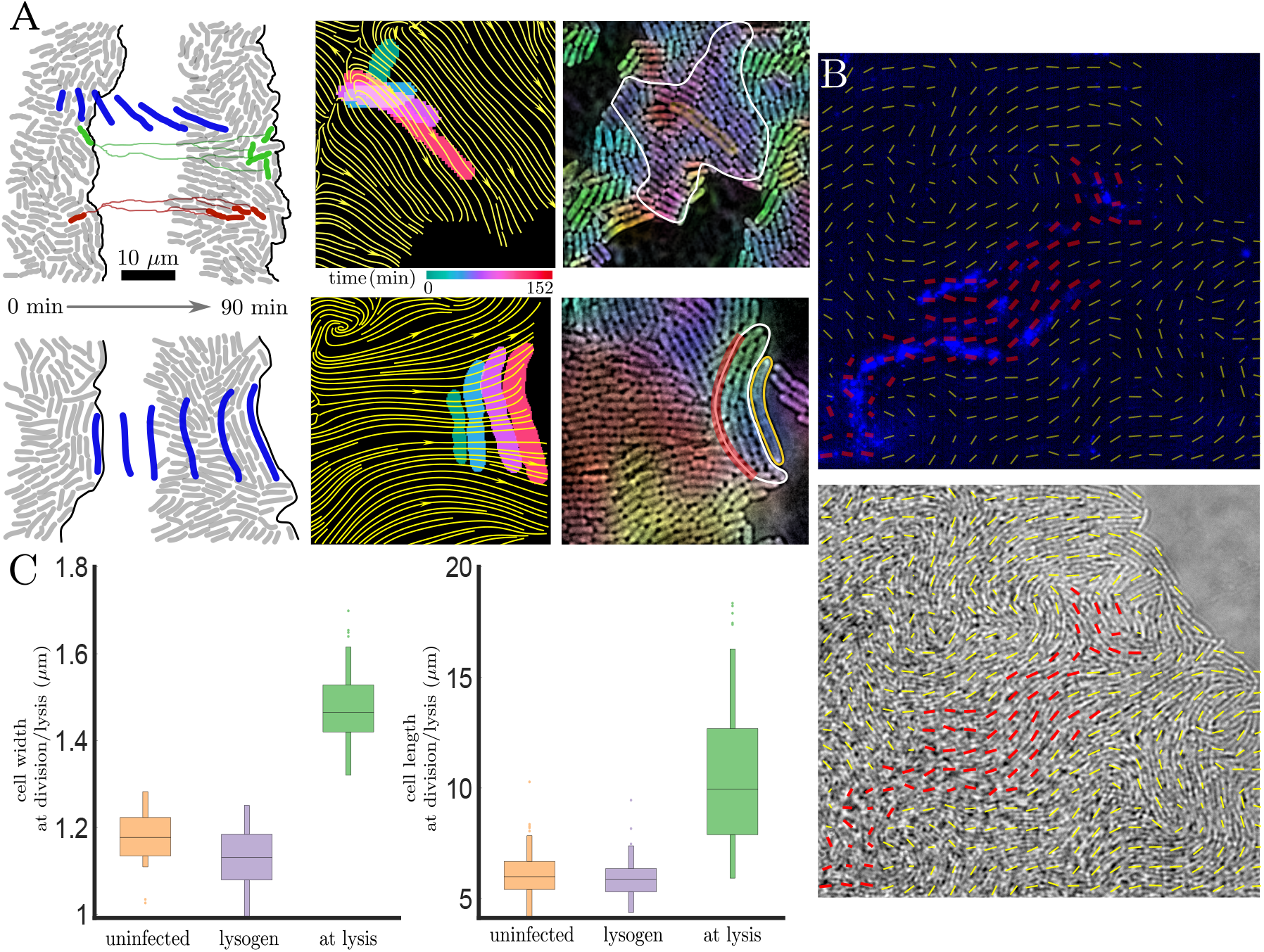
Bacterial advection, re-orientation. **A. (top)** A lytic cell just behind the edge of the colony is advected forward along with other cells, but is reoriented along the direction of the outward expanding growth front (indicated by yellow streamlines). **A. (bottom)** A lytic cell on the edge of the colony is able to temporarily surf the growth front by advection, remaining in parallel to the direction of the growth front (indicated by yellow streamlines). In both cases, the length of the lytic cell induces local alignment of neighbouring uninfected cells, orienting them parallel to the lytic cell. **B**. In the very early stages of the formation of a lytic river, the reorientation of the elongated lytic cell causes the local alignment of neighbouring cells, forming a nematic micro-domain. **C**. Lytic cells are unable to divide and continue to grow until lysis, elongating to 2.5x and widening to 1.4x an average uninfected cell.

This applies to uninfected cells. Following infection however, cells that enter the lytic pathway do not divide but elongate to around 2.5 times the length of an average uninfected cell (Burst length - 10.6 ± 3.1 *µm*, aspect ratio 8.5:1) (Figure 2**C**). This elongated length enhances the propensity of a lytic cell near the colony edge to reorient itself, like a wind vane, in the irection of colony growth [6, 35] (Figure 1**A**, colony growth streamlines marked in yellow). This results in lytic cells predominantly oriented along the direction of growth and the uninfected cells flow past these elongated cells due to their growth-driven advection, causing the lytic cells to be left slightly behind the propagating periphery.

### 2.3 Anisotropy of phage burst upon lysis

First we examined the expulsion of phages during the lytic event. We identified the point of release of phages from the lytic cells and mapped this onto a reference cell (Figure 3**B**) and plotted it as a heat map. The release of phages was found to be anisotropic, with a majority of bursts (*71%, 87 of 122 bursts*) occurring at the polar ends of the cell. No preference was found between the old and new poles. In conjunction with cell orientation data, near the edge of the growth front, phages are thus expelled predominantly parallel (in either direction) to the growth of the bacterial front. In addition to confirming this polar bias, SEM images also revealed that most often, the cell wall remained almost intact following lysis (Figure 3**A**).

**Figure 3:**
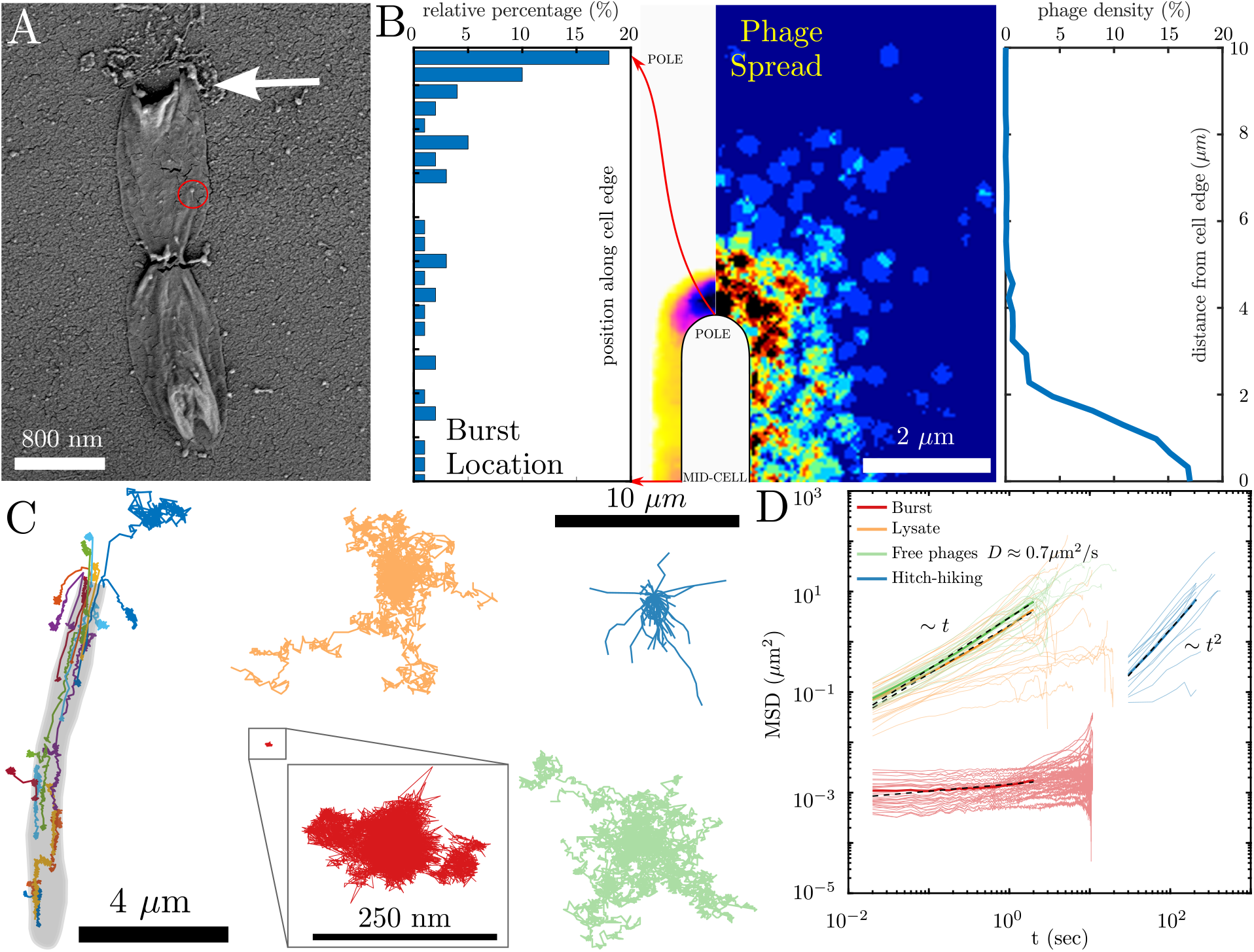
Phage localization, dissemination and diffusion. **A**. Anisotropic polar expulsion was confirmed by scanning electron micrograph (white arrow), also revealing the presence of an intact cell wall and individual phages (red circle). **B**. A heatmap of the positions of phage expulsion over a reference cell reveal a bias for the polar end *(71%)*, resulting in a polar bias of ejected phage particles, with almost *98%* of phages remaining within 4 *µm* of the cell. **C**. The lysis event was monitored by tracking individual phages inside the cell and following lysis (at 20 ms intervals), providing insights into the ejection, localization and diffusion of phages. **D**. Phages from unpurified cell lysate (orange) and purified cell lysate (green) displayed diffusive characteristics (MSD ~t, D *≈* 0.7 *µm*^2^/s), as illustrated by representative tracks. Ejected phages (Red) appeared to be caged in cell debris hindering diffusion. However, the hitchhiking phages (blue) were carried to much longer distances by cell advection, resembling ballistic motion (MSD *~t*^2^).

The spread of phages immediately following lysis was found to be localized, lying usually within one cell length (~ 4 *µ*m, Figure 3**B**, third panel), with the force of expulsion occasionally carrying them up to 10 *µm*. However, a significant phage density remained within the erstwhile cell, pointing to the hitherto unexplored role of cell debris in phage spread. With the cell membrane remaining following lysis, phages trapped within may not be able to infect any other cells, significantly reducing the effective burst number.

### 2.4 Phage diffusion and advection after a lytic burst

Tracking the positions of phages before and following expulsion (Figure 3**C**) revealed that the phages released were varied in their transport behaviour. While most phages remained close to the cell, potentially caged by cytoplasm and cell debris, only a few moved away (*<* 10%). Even among those that do, a range of movements from diffusive (diffusion constant *D ≈* 0.7 *µ*m^2^*/*s) to hindered diffusion are observed. Additionally, following cell lysis within a bacterial colony, diffusion was obstructed by the presence of neighbouring [19, 36] which further minimized the spread of infection within the colony.

In addition to phage spread by expulsion and diffusion, phages are also transported by hitch-hiking on bacterial cells [37]. Cell growth within a bacterial colony, while causing reorientation and alignment (Figure 2**A** blue cell) also generates an advective flow, pushing cells ahead in the direction of growth. This also carries along infected cells, allowing phages to travel with the advancing front. Crucially, near the growth front, cell advection is directional, radiating outward from the colony homeland. Hitch-hiking phages do not necessarily infect the specific bacterium on which they hike; on occasion, they may detach from the cell and freely diffuse before an eventual infection.

### 2.5 Phages surf the bacterial colony leading to the formation of ‘lytic infection rivers’

In our experiments, phage infections that remain confined within the homeland (outline marked in green dashes) form small ‘infection stars’ (Figure 1**A**) which are similar to plaques. Within the homeland, the cell density (40 to 50 cells/ 100 *µm*^2^) allows for little growth, whereas at the edge of the homeland, vast empty territory extends. Thus, infections at the edge of the homeland may exhibit a very different pattern, propagating with the expanding colony to form anisotropic ‘lytic rivers’ (Figure 1**B**) consisting of a core of lysogeny (Red) surrounded by a lytic layer (blue) which is able to spread the infection in the surrounding uninfected cells (Black). As we show below, cells within such lytic rivers exhibit an ordered orientation extending radially outward from the homeland, extending as far as the colony spreads. In comparison, the orientation of cells in infection stars and plaques is disordered.

A unique feature of the lytic river is the inward dip at the river mouth, formed by the relatively slower expansion of the front, presumably due to lytic death along the river. In the wake of the formation of the invagination, the lytic infection (in blue) now splits, infecting cells along both edges of the dip. The lysogenic cells that emerge occupy the empty space and, together, form the characteristic delta of the infection river. In the following sections, we examine the colonies at a smaller length scale to understanding the mechanisms that underlie formation and propagation of the surfing phage and lytic rivers.

### 2.6 Formation and growth of lytic rivers

Given these microscopic phage and bacterial growth and transport properties, we argue that lytic rivers form and propagate due to a combination of radially oriented elongated lytic cells, polar and localized cell phage expulsion and isotropic radial advection. The growth-driven reorientation of the lytic cells promotes the alignment of neighbouring cells, creating a nematic micro-domain of cells in the vicinity of a long lytic cell (Figure 2**B**). The phages that are then released upon lysis are either expelled beyond the growing bacterial front or are transported to the proximity of the front via a combination of diffusion and hitch-hiking. The front of an uninfected colony advances at an average of 1 *µm/min*, setting an upper bound to the advection of a cell, as the presence of the non-dividing, slower growing and periodically dying lytic cells, retards this progress. Although the diffusivity (Figure 3**D**) of unhindered free phages is sufficiently large to transport them over distances larger than the advective hitch-hiking transport on comparable timescales, the directed nature of advection is key for the formation of lytic rivers.

An important factor in infection spread is the growth phase of the bacterial cells, as the latent time of the infection is inversely proportional to the growth rate of the cell [38]. With only a narrow band of ~ 200 *µm* (Figure SI 2**C**) at the front of the expanding colony in a growing phase [9, 39], the spread of infection within the bulk, where most cells are in or approaching stationary phase, is arrested, remaining frozen in time. As a consequence, for the continued proliferation of a lytic river it is imperative that the infection is able to remain in that growing band, i.e., surf the colony front. This also restricts an existing lytic river from expanding laterally within the bulk of a colony (Figure SI 1**B**). The directed transport of phages radially outward ensures their continual presence proximal to the growth front of the bacterial colony.

The phages that surf the growth front can now diffuse and infect other cells at the very edge, ensuring that the infection persists at the edge. Thus, new infection streams are spawned, continuously distributed from the immediate vicinity of the lytic river to even over 100 *µm* away. When such an event happens relatively far away from the parent lytic river, it often gets extinguished (Figure SI 5**B**). However, events nearer the parent river can spread and merge into the parent river causing it to expand outward.

Generally, a growing *E. coli* colony of uninfected cells expands radially, maintaining a circular shape. This is also true for a mixed colony of uninfected and lysogenic cells. With both having almost identical growth characteristics (Figure SI 2**A**), neither has an advantage on range expansion. Cell lineages of both populations form sectors [5], which compete to expand the growing front of the colony (Figure SI 6, simulation). However, with lytic cells in the mix, with their slower growth, elongated length and, crucially, now cell death in the colony, it changes the dynamics of colony growth and the relative fraction of the different cell types.

A key difference is that death only occurs within the uninfected population and not the ‘resistant’ lysogenic population, thereby offering the lysogens an advantage in spreading.

Consequently, the ratio between these species in the *initial* seed population plays a significant role in determining the infection pattern outcomes (Figure 1**B**). In a seed of bacteria and phages with a low average phage input (API, ratio of phage to bacteria), lysogenic cells are unlikely to emerge. The dominant uninfected cell population can quickly smother any lytic infection, and expand circularly, similar to a single species colony. At a high API, infection outcomes are biased to lysogeny [29, 30], which now behaves as a phage absorbing barrier. The colony grows out circularly, similar to a mixed colony, with uninfected cell sectors that are often protected by neighbouring lysogenic cell sectors. However, at an intermediate range of APIs, lytic cells are found in relative higher numbers. This allows more infection rivers to grow from the homeland and compete to surf the front of the expanding colony, often coexisting with uninfected and lysogenic cells.

Note that the dynamics at the scale of a single lytic river is not determined directly by the API. That is, if the same local arrangement of lysogens, uninfected bacteria and phage happened to arise in some portion of two different colonies with different APIs, the subsequent stochastic dynamics in those local regions would be the same. Instead, the reason very different outcomes are seen for different API is that it determines the number of infections that are initially near the boundary of the homeland and thereby the number of infection rivers that grow outward. At very low APIs, there are no infections at the edge and one only sees infection stars within the homeland. As the API increases, relatively isolated rivers form and can grow substantially along with the lysogenic cells within them. Eventually, rivers will merge with neighbouring ones and only lysogens will be left at the growing front. At even higher APIs, there are many initial infections on the edge of the homeland and lysogeny is favoured due to the high multiplicity of infection [31, 40, 41]. This inhibits the expansion of the lytic river and allows uninfected cells to outgrow them. The result is a coexistence of sectors consisting of uninfected and lysogenic cells. Thus, the growing colony effectively retains long-term memory of the initial condition, i.e., the spatial arrangement of lysogens, uninfected bacteria and phages inside the homeland area.

### 2.7 Lattice model incorporating advection by cellular growth

Finally, we constructed a simple lattice model to determine the minimum ingredients required to reproduce the observed pattern. The model is defined on a two-dimensional square lattice. Each lattice site can be occupied by at most one bacterium, but can contain multiple phage particles. The important features of the model are: (i) the bacterium division rate at a site is an increasing function of the empty lattice sites within a radius of 100 sites around the site, (ii) advection of the bacteria via division of the cells inside a colony is taken into account by allowing chains of cells to be pushed outward upon a division, (iii) phage particles diffuse rapidly, but adsorb to a cell relatively fast if they reach a site that contains a bacterium, (iv) an infected bacterium chooses lysogenization if it is infected by a second phage within short time interval, and otherwise chooses lysis, reflecting the MOI dependent decision making, (v) upon lysis, the phages are placed at the burst site and the cell is removed. The detailed model description and parameters are given in the Methods section. Note that the model considers neither the anisotropy due to the elongated shape of the cells, nor the extra elongation of lysing cells.

We start each simulation with a circular region in the centre of the lattice partially occupied by cells. In the following, the unit time is set so that the maximum possible growth rate in the model is 1. When there are no infected cells or phages in the system, after about time *t ≈* 12, the system reaches a state where a roughly circular colony radius expands at a constant speed of ~ 30 lattice sites per unit time.

Figure 5 show the resulting colonies with various values of *API* and free phage hopping rate *p*_*diff*_ (proportional to the phage diffusion constant). For small *API* values, we observe that only a few of the lysis events managed to reach the edge of the home land. The subsequent lysis events propel some phage particles outside of the colony edge, which then diffuse quite far along the edge, starting new lytic rivers. Note that the advection of cells by growth is essential to get the elongated lytic rivers in radial direction (see Figure SI 6 for comparison with the simulation with very limited advection).

Merging of the lytic rivers results in the widening of the lysogen region within the lytic rivers. The number of the lytic rivers formed outside of the homeland increase with the *API*. This naturally results in the lysogeny dominant pattern for high *API* case, while uninfected cells dominate the pattern in the low *API* case.

The frequency of starting new lytic rivers as well as merging of lytic rivers strongly depends on the phage’s ability to spread by diffusion at the edge of the colony. When *p*_*diff*_ is small, the expansion of the lysogenic region relies on the merging of existing lytic rivers, hence it happens only when *API* is high. For high *p*_*diff*_, new lytic rivers are continuously formed at the edge of the colony around the existing lyric rivers, resulting in expansion of the lysogenic region from initially a single lytic river.

It is worth noting that the how much phage diffuse in the colony before adsorption is an important parameter. In the bottom row of Figure 5, the phage adsorption rate *η* is increased by 10-fold compared to the rest. When a bacterium bursts in the colony, most of the phages are quickly adsorbed before diffusing far, resulting in high MOI infections in the neighbors and hence, more formation of lysogens and less lysis. Therefore, most of the lytic rivers die out rather quickly. These effects are quantified in (Figure SI 7) via the spatial distribution of the infections with *MOI* = 1 and higher *MOI* infection per burst.

## Conclusions and Discussion

We have explored the interplay between phage infection and bacterial growth on an agar plate. We find that a growing colony of *E. coli* infected with the temperate phage *λ* exhibits intricate spatio-temporal dynamics that depends on the initial ratio of phage to bacteria (API). Over large length- and time-scales, we find a variety of outcomes: as the API increases from low to high, we see dominance of uninfected bacteria due to the quick extinction of phage, then emergence of lytic rivers enclosing lysogen sectors that grow and eventually dominate the colony, followed by lytic rivers that get extinguished, causing the dominance of uninfected bacteria, and finally lytic rivers that get extinguished but lead to co-existence of lysogens and uninfected bacteria in adjacent sectors. It is interesting that lytic events do not spread in the bulk of the colony but remain confined in these “rivers” which grow roughly radially outward along with the growing colony.

This long length- and time-scale behaviour emerges from the interplay between mechanics and growth and infection at small (cellular) length scales and short (one cell generation) timescales. We showed that mechanical forces coupled with cell growth result in local nematic ordering of cells in the colony – cells near the growing front are oriented perpendicular to the growth direction, while cells further in but still near the front are oriented parallel to the growth direction. Cells that have been infected by the phage and entered the lytic pathway are elongated and the mechanical forces preferentially orient them parallel to the growth direction. This, combined with the polar release of phage upon a lytic event and radial advection of phage that are adsorbed to the surface of bacteria, results in the lytic rivers growing primarily in the radial direction at the growth front. Lateral growth of the rivers occurs at a slower pace due to diffusion of phage at or just beyond the growth front, resulting in nucleation of new lytic rivers nearby that merge with the parent river, producing the characteristic delta shape as it grows.

To better understand how the long-term coexistence of uninfected cells, lysogens and lytic bursts across the colony, or lack of such co-existence, emerge from small length- and time-scale processes, we constructed a lattice model of growing bacteria infected by temperate phage. An important feature of our model is that it explicitly includes the growth-driven advection of bacteria and of phage adsorbed to these bacteria. We implement this using an algorithmic rule that results in stochastic “pushing” of bacteria radially outwards when a bacterium within the bulk of the colony divides. Such growth-driven advection arises in models that represent individual bacteria and all the forces on them explicitly [20], and was also implemented in coarse-grained lattice model that treat biomass as a continuous variable as cell-shoving [42]. In our context, the former type of model is computationally intractable for the colony sizes we are examining, therefore it was important to implement a coarse-grained rule for bacterial advection in a lattice model. Furthermore, we limited the bacterium cell per lattice site to be at most 1, which allowed us tracking each cell’s fate including MOI with a rather simple algorithm. The model captured the basic pattern where uninfected bacteria dominate at low API, but at high API they co-exist with lysogens each in their separate growing sectors. The model also allows us to understand how this pattern depends on parameters such as phage diffusivity, rate of advection, etc. Thereby, we can show that both advection and hitchiking of phage on advecting bacteria is necessary for the formation of lytic rivers and the patterns of co-existence/dominance as a function of API.

The model does not incorporate anisotropy of the lytic burst or nematic ordering of uninfected and infected cells. One difference between the model and experiments is that in the model the edges of lytic rivers grow radially on average. Whereas, in the experiments, for API values just above the lowest we examined, we see a clear outward curvature of the edges of the rivers (see Figure 4**B**) which results in lysogens eventually dominating the colony. This is not observed in the model, because we speculate the anisotopy of the burst combined with the nematic ordering of the cells are necessary for this phenomenon which was not factored in. In the experiments we also observe that the lysogenic region within the delta of a lytic river lags a little behind the growing uninfected front, although the uninfected and the lysogenic cells grow at the same rate (Figure SI 3). This happens because within the lytic river, the lytic cells cease to divide and eventually lyse, hindering the growth of the front. This could be exacerbated by time taken for a multiply infected cell to turn lysogenic and resume growth.

**Figure 4:**
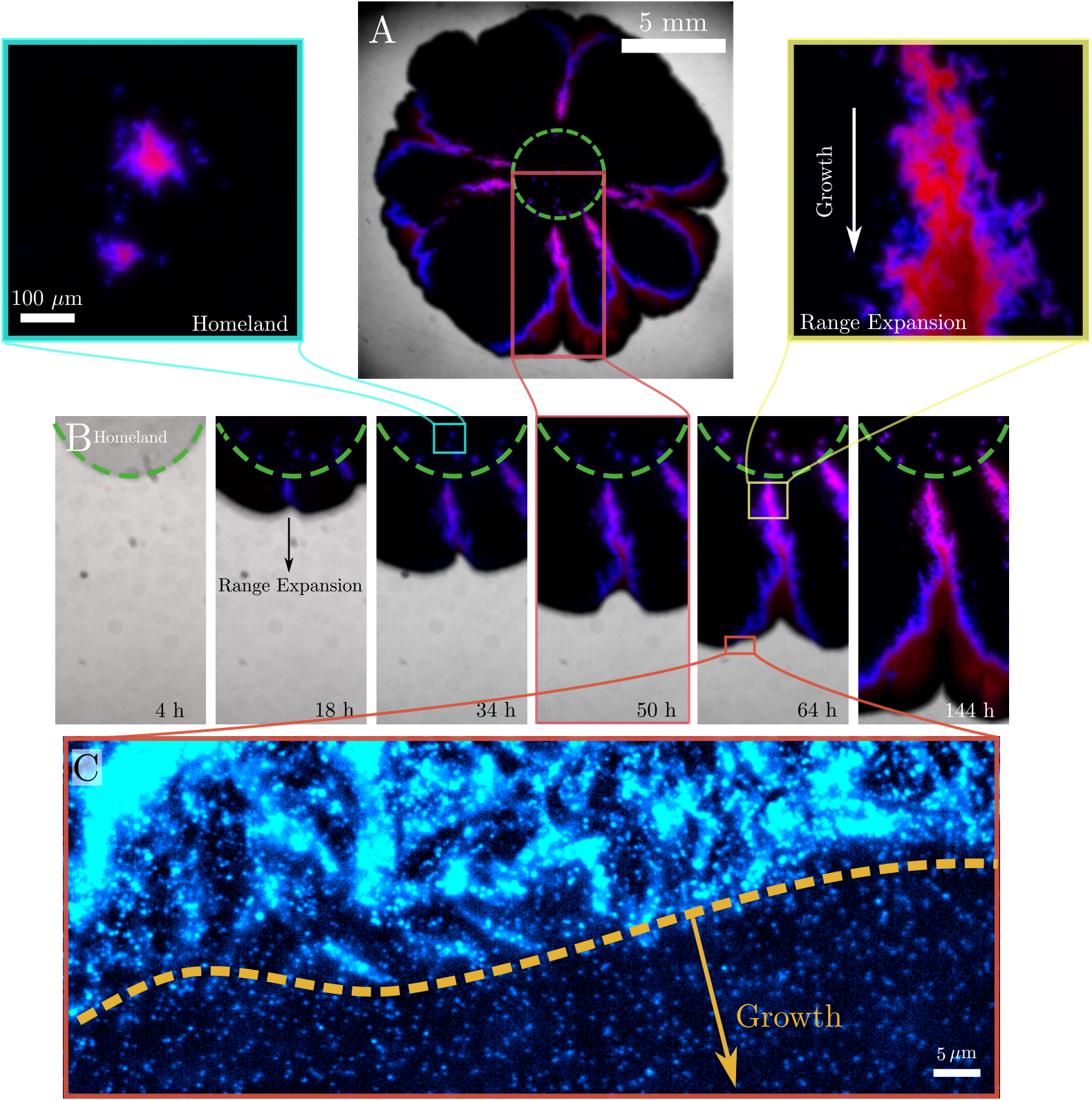
Infection dynamics and lytic rivers. **A**. The homeland (bound by green dashes) contained an initial seed of phages (~ 10 phages) mixed with uninfected cells (~ 10^5^ cells). Infections within the denser constrained homeland form distinct lytic ponds, while infections at the edge of the homeland can propagate radially outward as lytic rivers. **B**. A lytic river propagates slower than uninfected regions, resulting from the switching of dividing uninfected cells (Black) to non-dividing lytic cells that eventually die. These form the bounds of the lytic river (blue), allowing only resistant lysogenic cells (Red) to grow within. **C**. Some lytic rivers may die out, however the presence of infected cells and phages at the edge of the growth front ensure continued infection.

**Figure 5:**
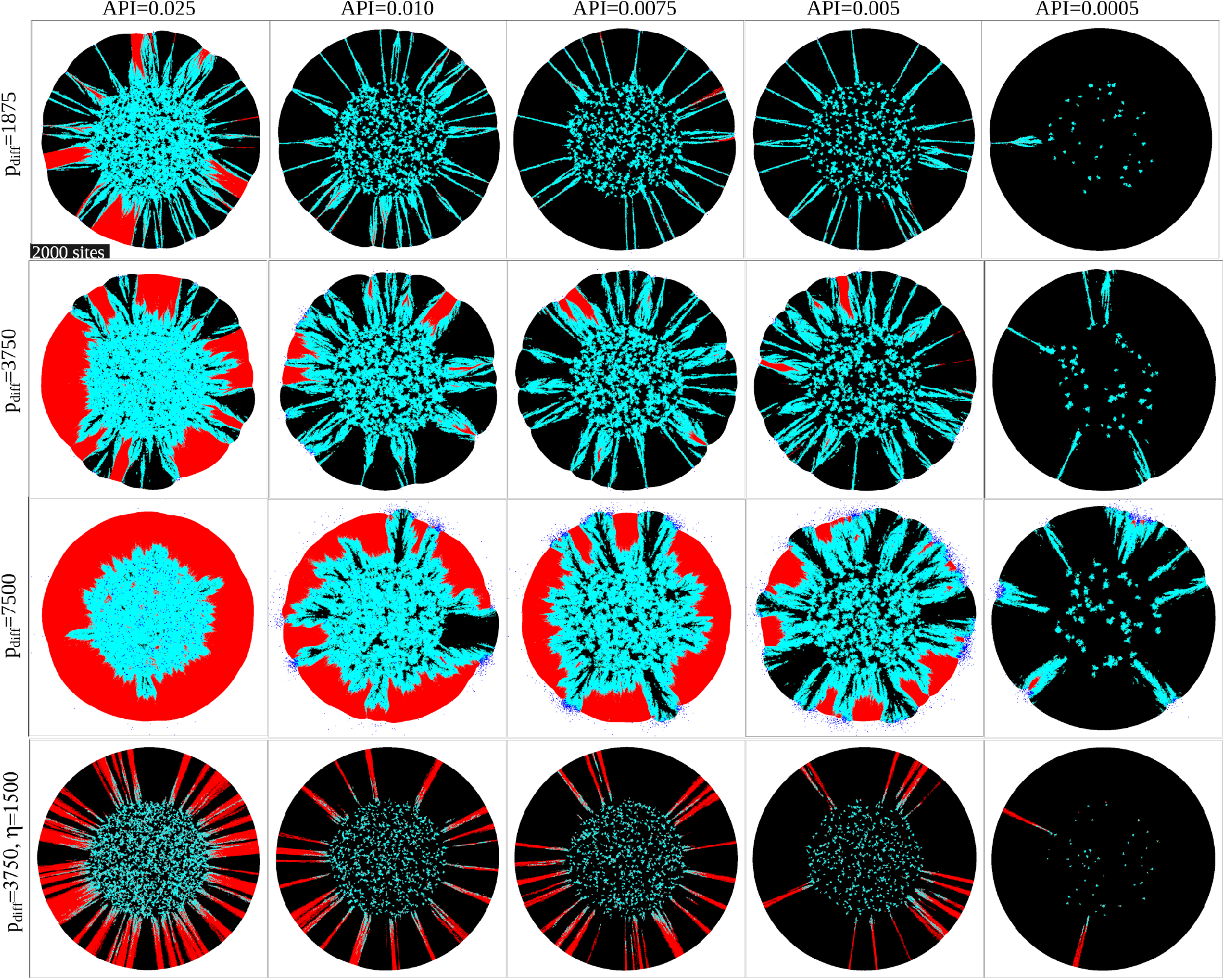
Model results. Black is an uninfected cell, yellow is a just infected, undecided cell, red is a lysogen, cyan is a cell in the lysis pathway. Dark blue sites are the sites that contain free phages (shown bigger for visual clarity). All the images are the snapshot at time 54. The value of the phage diffusion parameter *p*_*diff*_ is increased from top row to the third row. In the bottom row, the *p*_*diff*_ is set to the middle value and the phage adsorption rate *η* is increased by 10-fold, resulting in much less diffusion of phages in the colony before adsorption.

This lag of the lysogenic front in some cases allows nearby uninfected bacteria to grow into the space in front of the lysogens, thereby cutting them off and extinguishing the lytic river and resulting in a rougher colony boundary (Figure SI 4). In our model, lytic rivers instead get extinguished purely due to the random meandering of their boundaries, and colony boundaries are therefore smoother. Despite these caveats, our model provides strong evidence for the importance of advection and phage hitchiking in producing lytic rivers, and can be extended in future to account for other processes, such as anisotropy of the phage burst.

The emergence of large scale features (coexistence vs. dominance; confinement of lytic events in radial rivers) from the interplay of mechanical forces, bacterial growth and phage infection processes at small scales is interesting in itself, but it also has implications for the evolution of bacteria-phage interactions. Although we have not observed the growing colonies for long enough for mutants to emerge, the timescales over which evolutionary changes occur in phage and bacteria is not much larger than the timescales over which these colonies grow. At both low and high API where lytic events rapidly disappear, there is not much scope for the evolution of phage. However, at intermediate API values (Figure 1**B**) where lytic rivers persist for a long time, one can speculate that a co-evolutionary arms race between the phage and bacteria will occur. Phage mutations that increase their infectivity will result in the lytic river edge curving outward faster, whereas mutants with lower infectivity will probably be left behind by the curving river comprised of the original phage. Similarly, bacteria that become more resistant to phage infection would cause the river edge to curve inward. Thus, just as the movement of sector boundaries in purely bacterial colonies signal the emergence of mutants [8], the meanderings of observed lytic rivers could indicate the progress of the phage-bacterial arms race. The evolution of the lysis vs. lysogeny decision and its dependence on MOI is also interesting factor to consider in this spatially structured expansion. *λ* phage is known to go lytic upon single infection and tend to go lysogenic upon double (or more) infection. By analyzing the population dynamics in a well-mixed system, lysogenization has been speculated to be a bet-hedging behavior to survive in fluctuating environment where phage and host have different survival probability [43, 44], where MOI contains the information of the remaining hosts in the environment [41]. Another work showed that it is also beneficial for phages competing to invade a large population of sensitive hosts [40]. However, the benefit of MOI dependent decision is less clear in the expanding colony where the cells are locally densely packed, because high MOI infection is likely to happen almost always around the phage burst and the occurrence of low MOI events relies on some phages happen to escape to reach hosts further away. Hence, from the perspective of phage expansion which requires lysis to keep happening, it is not trivial if the MOI dependent decision is beneficial. Any mutation in this “rule” would likely show up in the size of the lytic river delta, how rapidly a newly nucleated river splits, as well as how long the river survives. The co-evolution in the current setup can be a relatively easy way to test if the “rule” the phage has employed is robustly beneficial in the spatially structured system.

## 3 Methods and Materials

### 3.1 Experimental Details

The phage-*λ* virus has a temperate life cycle that allows it to either (i) reside within the genome of *E coli* through “lysogeny” (making the bacterium heritably resistant to further viral infection) or (ii) enter into a “lytic” phase, during which it kills and lyses the cell to produce offspring (the viral numbers increase multi-fold). Of particular note here is that the infection outcome *i*.*e*. lysis or lysogeny, depends on multiple intrinsic and extrinsic factors (*e*.*g*. environmental, bacterial densities, growth rates etc.) and one of the more important factors is the so called multiplicity of infection (MOI) – if a single bacterium is infected, within a certain time window, by two or more viruses, then the bacterium enters lysogeny, and lysis otherwise. The experimental system that we use [29] allows us to visualise these different infection states: briefly, the phage capsid decorative protein, gpD is translationally fused with the fluorescent mTurquoise2 (blue) molecule so that single phages can be visualised. These phages are therefore visible before infection, as bright spots, and upon lytic decisions the cells produce progeny phages and the cells themselves appear full of blue spots in lytic state. Lysogenic reporters are constructed by transcriptionally fusing mKate2 (Red) to the phage lytic repressor gene, cI. Upon lysogenic decisions, cells express the lysogenic reporter colour as they grow and divide.

#### 3.1.1 Strains and growth conditions

All experiments were performed using lysogeny broth (10 gL^*−*1^ Tryptone, 5 gL^*−*1^ yeast extract, 5 gL^*−*1^ NaCl) supplemented with 10 mM MgSO_4_ and 0.2% maltose, hereby referred to as LBMM. Phages were extracted from lysogenic LZ1367 *E coli* (MG1655 (*λ*D-mTurquoise2 cI857-mKO2 bor::CmR)[pACYC177-PLate*D] obtained from Lanying Zeng) as reported previously [45], yielding a phage stock of 10^11^ phages/mL in SM buffer. Briefly, a 100 mL culture of lysogenic LZ1367 was grown to an OD just under 0.4 in LBMM at 30 °C. It was induced at 42 °C for 20 min and then moved to 37 °C until clear (~ 1 h). Following addition of 5 mL of chloroform, the lysate was incubated at 30 °C for 30 min. Debris were removed by centrifugation at 10K rpm for 20 min at 4 °C, after which 5.84 g NaCl was added to the supernatant and cooled on ice for 3 h. The phage suspension was purified by centrifugation again and 10 g of PEG 8000 was dissolved and left on ice overnight. The following day, following a third centrifugation, the supernatant was discarded and the PEG 8000 pellet was dispersed in 1.5 mL of SM buffer. The PEG 8000 was removed by three cycles of the addition of 2 mL of chloroform and centrifugation at 3K rpm for 20 min at 4 °C to obtain the phage stock.

*E coli* MG1655 wt was used in experiments and following infection the lysogenized variant was isolated. However, the *E coli* used in this study did not contain the plasmid pACYC177-PLate*D (for the production of the wt gpD capsid proteins) and hence the capsid of phages produced from lytic infection and induction of lysogenic cells were made exclusively of the fusion protein.

#### 3.1.2 Colony-level infection spread experiments

To study the spread of phages in a growing *E coli* colony, 50 *µ*L of an overnight cell culture was inoculated into 5 mL LBMM and grown for 3 h at 30 °C. To 100 *µ*L of cell culture that was diluted ten times (10^8^ cells/mL), 1 *µ*L of phage at different concentrations were added, resulting in APIs ranging from 1 to 10^*−*4^. The mixture was kept for 15 min at 4 °C to allow for phage adsorption and then for 15 min at 30 °C before a 1 *µ*L spot was placed on a 1.7% LBMM agar plate, which was kept in an incubator at 30 °C to grow. Using a Leica M205 FA stereo microscope, brightfield and fluorescence images (GFP channel for phages – Ex: 470/40 and Em: 525/50 and RFP channel for lysogeny – Ex: 546/10 and Em: 605/70) were captured either every 24 h or for time-lapse imaging, every 20 min for several days. During time-lapse imaging, temperature was maintained by a placing the agar plate on a custom-made temperature-controlled ITO plate.

### Microcolony-level infection spread experiments

In order to study individual cells in a growing microcolony, we devised a new set-up to allow visualisation under an inverted microscope. Cells were imaged every 4 min using an Olympus IX81 inverted microscope with and attached aCMOS camera (Photometerics Prime). A coolLed pE 4000 lamp was used as a light source for fluorescence imaging, specifically Ex: 435 nm and Em: 458/480 nm for phages and Ex: 550 nm and Em: 555/58 nm for lysogens. Temperature was maintained by a stage-top incubator (Okolab). To briefly describe the agar platform - a hole was punched in the base of a 35 mm petri dish, and from below a glass coverslip was taped on. LBMM agar was then poured into the dish and cooled. The glass cover was removed, cells were placed on the agar and the glass cover replaced to allow visualization under the microscope. This also minimized evaporation of the agar, enabling longer experiments. To study the spread of infection in a microcolony, we required a single lytic cell within or neighbouring a microcolony of uninfected cells. To achieve this, we mixed uninfected cell in exponential phase with lysogenic cells in the stationary phase (approximately 4:1). This mixture was diluted and a drop was placed on agar, where it grew for 40 min at 30 °C. During this time the uninfected cells multiplied, whereas the lysogenic cell was only emerging from lag. The petri dish was then transferred to 42 °C for 20 min for induction and then back to 30 °C for imaging.

### Mapping the position of phage release and phage spread

Lysogenic cells were grown for 2.5 h at 30 °C following which the centrifuge tube was moved to 42 °C for 20 min for lytic induction and then back to 30 °C for 20 min for phage maturation. 1 µL of the cell suspension was sandwiched between two pre-warmed coverslips, which was placed on a microscope stage-top incubator maintained at 30 °C for imaging. Phage distributions were captured in the CFP channel using a 100x oil immersion objective at 4 s intervals. For each cell, the position of phages prior to and after lysis was captured, and the point of release of phages from the cell was marked on a reference cell. Using ImageJ, the images were binarized and employing a custom written MATLAB code, heatmaps were generated for the location of point of phage release and the distribution of phages. Additionally, the lysis event was also captured at a high time-resolution of 50 fps. However, this most often resulted in bleaching of phages and so was not extensively employed.

### Diffusion measurements

Diffusion of phages was measured in 2D on agar by tracking the movement of phages at 50 fps. Phage positions were obtained using the Mosaic particle tracker plugin in ImageJ [46], which were then used to construct MSD plots and obtain the diffusion coefficient. Positions were obtained for phages extracted from induced lysogenic, both with and without purification by centrifugation. Additionally, we also tracked phages that were expelled from cells that grew and lysed on agar. Purified phages were diffusive with a diffusion coefficient of 0.7 *µ*m^2^*/*s, which is of similar order to the diffusion coefficient for the *λ* phage found in literature [47].

## 4 Model

### Model description

Our model is defined on two-dimensional square lattice with linear size *L*, in total *N* = *L*^2^ sites. Each lattice site can be occupied at most one bacterium, while multiple phage particles can be located on one lattice site.

We assume that the growth of bacteria, as well as the development of phage in an infected cell, is affected by the availability of nearby space, mimicking inhibition of the growth by limitation of physical space as well as the limitation in nutrient availability. More precisely, we keep track of the fraction *f*_*i*_ of the unoccupied lattice sites within the Euclidean distance *r*_*f*_ of a given lattice site *i*. We assume that the growth rate is proportional to the following function:

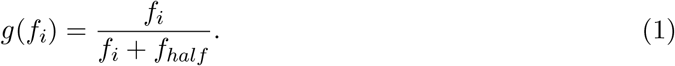

We assume that *g*_0_*g*(*f*_*i*_) is the growth rate of the bacterium at the lattice *i*. The half-speed empty site fraction is given by *f*_*half*_. Because 0 *≤ f*_*i*_ *≤* 1, the maximum growth rate is *g*_0_*/*(1 + *f*_*half*_). The division of the bacterium is modelled as *M*-step sequential reaction step, i.e., the bacterium at site *i* in state *k* becomes state *k* + 1 at the rate *g*_*M*_ = *M · g*_0_*g*(*f*_*i*_). *k* = 1 for the just divided bacterium. When the state *k* reaches *M*, the cell divides and a new cell is formed, and both the mother cell and the progeny cell goes to the state *k* = 1. This makes the average generation time for bacteria to be *M/g*_*M*_.

We model the advection caused by cell growth by letting a newly formed cell “push” one of its neighbour cell out towards the closet empty site. For example, if a cell at the coordinate (*i*_*x*_, *i*_*y*_) divides, and if it’s closest empty site (if there are multiple empty sites with the equal distance, one of them is chosen randomly) is at (*j*_*x*_, *j*_*y*_), then the pushing by one lattice site happens towards *α*-direction with probability |*j*_*α*_ *− i*_*α*_|*/*Σ_*α*_ |*j*_*α*_ *− i*_*α*_|, where the direction is positive (negative) if *j*_*α*_ *− i*_*α*_ is positive (negative). When a cell is pushed by one site, the free phage at the same site is also pushed with it. This continues until a cell is pushed out to (*j*_*x*_, *j*_*y*_) and there is no overlapping of cell.

The phage particles diffuses on the lattice, and each particle infect a bacterium in the same lattice site at a constant rate *η*, if there is a bacterium at the site. Infected bacterium will first go into the undecided infected state. If another phage infect such a bacterium before the commitment to the lysis is fully made, the bacterium becomes a lysogen. A undecided infected cell commit to lysis at a rate *p*_*dec*_ = *r*_*dec*_ *· g*(*f*_*i*_) + *ϵ*. Here, *r*_*dec*_ characterize the decision making time scale. This is to reflect the observed multiplicity of infection (MOI) dependence of the lysis-lysogeny decision making. We assume that the decision making slow down as the bacterial growth slow down due to lack of nutrient.

When the cell is commited to lysis, after some time the cell burst and the *N*_*b*_ phage particles are placed at the burst site the bacteirum is removed. The lysis development process is modelled as (*M−*1)-step sequential reaction step. The reaction rate at the site *i* for the phage development process is given by 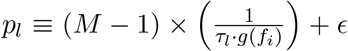 where the *τ*_*l*_ characterizes the phage latency time, which is assumed to be proportional to bacterium growth rate. When *ϵ* is non-zero, phage development can slowly happen at the rate *ϵ* even if the growth of the bacterium is completely halted. The lysogen is chosen only if the second phage infect the bacterium when it is still in the just infected state.

When a infected cell reaches the *M −* 1-th state, the cell burst and produces *N*_*β*_ phage particles. These particles are distributed in space following the Gaussian distribution centreed at the bust site with the standard distribution 1 lattice constant. The free phages will move to one of it’s 4 nearest neighbor lattice site at a rate *p*_*diff*_, allowing them to diffuse.

These rules are applied at a constant time step *dt*. For each time step, the following procedures (1-4) are repeated for *N* times:

1. Choose a random lattice site on the lattice.
2. **Update the bacteria cell growth**
3. If the chosen site contains a sensitive bacterium or a lysogenic bacterium, the cell growth state proceed to the next state with probability *g*_*M*_ *· dt*. If the cell division happens, pushing process as described is also executed.
4. **Update the bacterium status in undecided infected state and in the lytic pathway**
5. If the chosen site contains a bacterium in undecided infected state, the commitment to lysis is made with the probability *p*_*dec*_ *· dt*. If the chosen site contains a bacterium in the lytic pathway, the status of the cell is moved to the next state of lysis process at the probability *p*_*l*_ *· dt*. If the burst happens, the *N*_*b*_ phages are placed at the site and the bacteium is removed from the site.
6. **Update the phage adsorption and diffusion**
  a. If the chosen site contains phage particles, then the following procedure is done for each phage particles. If there is a bacterium at the same site, at the probability *η · dt*, the phage infect the overlapping bacterium. If that bacterium was uninfected, the bacterium’s state is updated to undecided infected state, with their growth state information preserved. If the bacterium was in undecided infected state, the bacterium’s status is updated to lysogenic of the given growth state. If the infection does not happen, the phage will move to randomly chosen next nearest neighbor site at probability (*p*_*diff*_ *−η*) *· dt*.
  b. If there is no bacterium at the same site, will move to randomly chosen next nearest neighbor site at probability *p*_*diff*_ *· dt*.

### Parameters

We set *g*_0_*/*(1 + *f*_*half*_) = 1, i.e., the unit time was set so that the maximum growth rate is one. We set *f*_*half*_ = 0.5. The length scale *r*_*f*_ to check the neighbor empty sites was set to 99.5, to be consistent with the observation that the bacteria is metabolically active within about 200 *µ*m from the edge of the colony, and that the linear size of a bacterium is about 1 *µ*m. The number of growth steps is set to *M* = 10, giving about 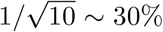 variation in the division time. The lysis-lysogeny decision rate *r*_*dec*_ is set to 15, and the lysis latency time *τ*_*l*_ is set to 2.2.

Three different phage hopping rates were tried to demonstrate the dependence of the pattern on the parameter: *p*_*diff*_ = 1875, 3750, and 7500 (Assuming lattice site linear length is 1 *µ*m and the maximum division time to be 40 min, these values correspond to the phage diffusion constant ranging from 0.2 to 0.8 *µm*^2^/s) phage infection rate to an overlapping bacterium is set to *η* = 150 unless otherwise noted, and the burst size *N*_*b*_ = 10. *ϵ*, the rate at which non-growing infected bacterium processes the phage decision and the burst, was set to 0.015.

The time step *dt* was set to 1*/p*_*diff*_, since *p*_*diff*_ is the largest rate among parameters in the model. The system size *L* was set to be large enough so that no bacterium or phage reaches the edge of the system with in the simulated time. The presented results used *L* = 6000.

### Initial condition

To mimic the initial inoculation of the cells on agar pad, randomly chosen 1.4% of the sites within the Euclidean distance of 1500 from the centre of the system was occupied by bacteria. This constitutes the “homeland” in the simulation. We vary the initial average phage input *API* as a parameter, and according to the Poisson statistics, a bacterium was kept uninfected with probability *e*^*−AP I*^, while assumed to be infected by only one phage and and to be in the lysis pathway with the probability *API·e*^*−AP I*^. With the probability (1*−e*^*−AP I*^ *−API·e*^*−AP I*^), which is the probability to be infected by two or more phages, the bacterium was set to be a lysogen. The growth state (the phage development) state of a uninfected (infected) bacterium is chosen randomly out of the *M* (*M −* 1) possible states.

## Acknowledgements

We acknowledge support from the Department of Atomic Energy (India), under project no. RTI4006, and the Simons Foundation (Grant No. 287975). ST acknowledges funding from the Human Frontier Science Program and the Max Planck Society through a Max-Planck-Partner-Group.

S.K. also acknowledges funding from the Science & Engineering Research Board, Department of Science & Technology, Government of India (Matrics grant MTR/2020/000253). N.M. acknowledges funding from Novo Nordisk foundation (Grant Ref. No. NNF21OC0068775).

## Supplementary Information

**Figure S1:**
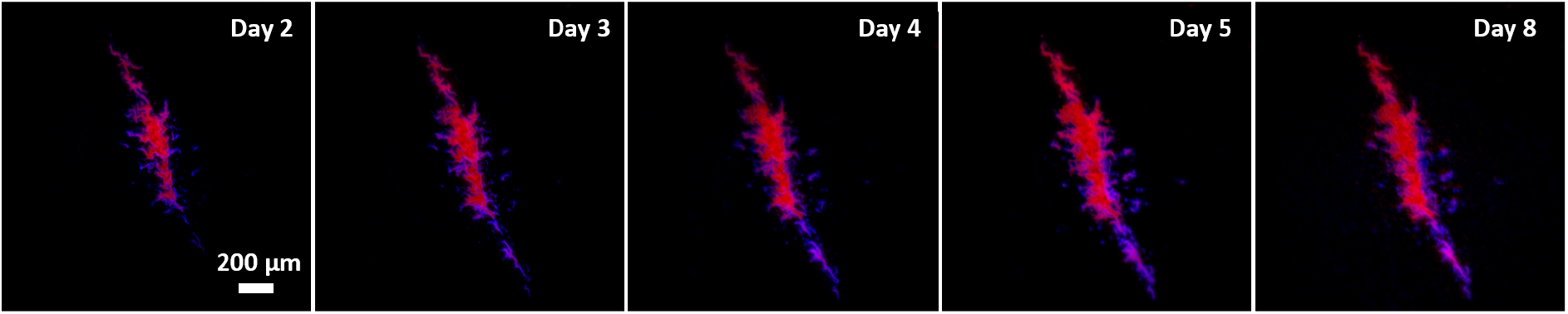
Anisotropic spread of infection within the colony. The lytic river grows in the direction of colony growth (Direction of colony growth from top to bottom)

**Figure S2:**
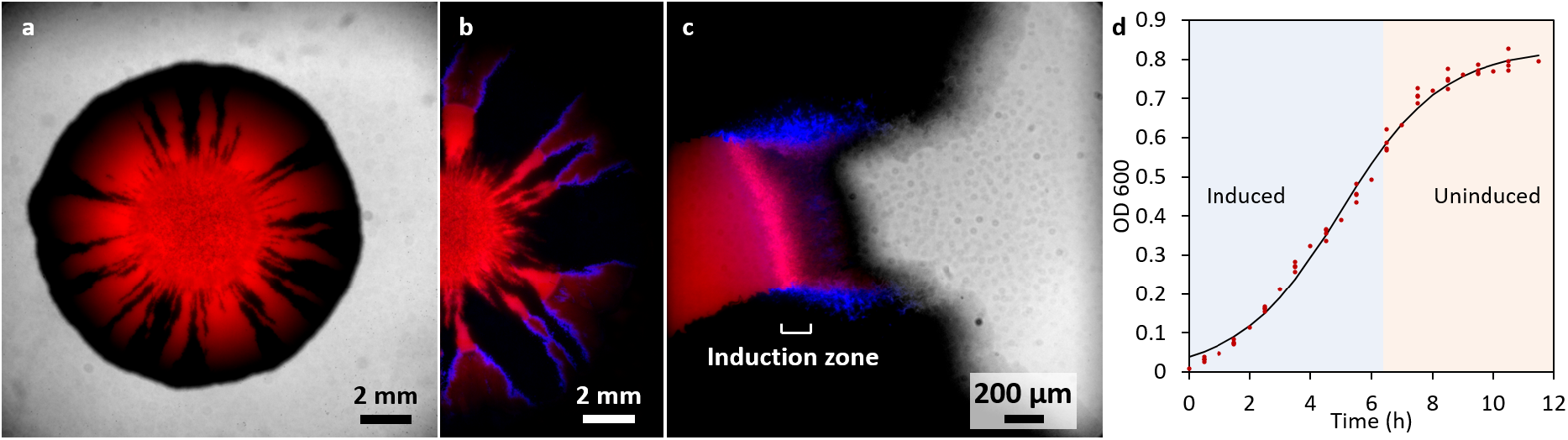
Measuring the width of the induction zone at the colony front. **A**. A colony of wild-type (Black) and lysogenic (Red) *E*.*coli* in equal proportions imaged in brightfield and mRFP channel. **B**. Induced colony after 12 hr displaying lysis along the boundary of the *wt* and *ly* cells. **C**. Induction zone was measured to be ~200 *µ*m in width as observed by the increased fluorescence in CFP. **D**. Growth graph demarcating the induced and uninduced phase of the lysogenic cells.

**Figure S3:**
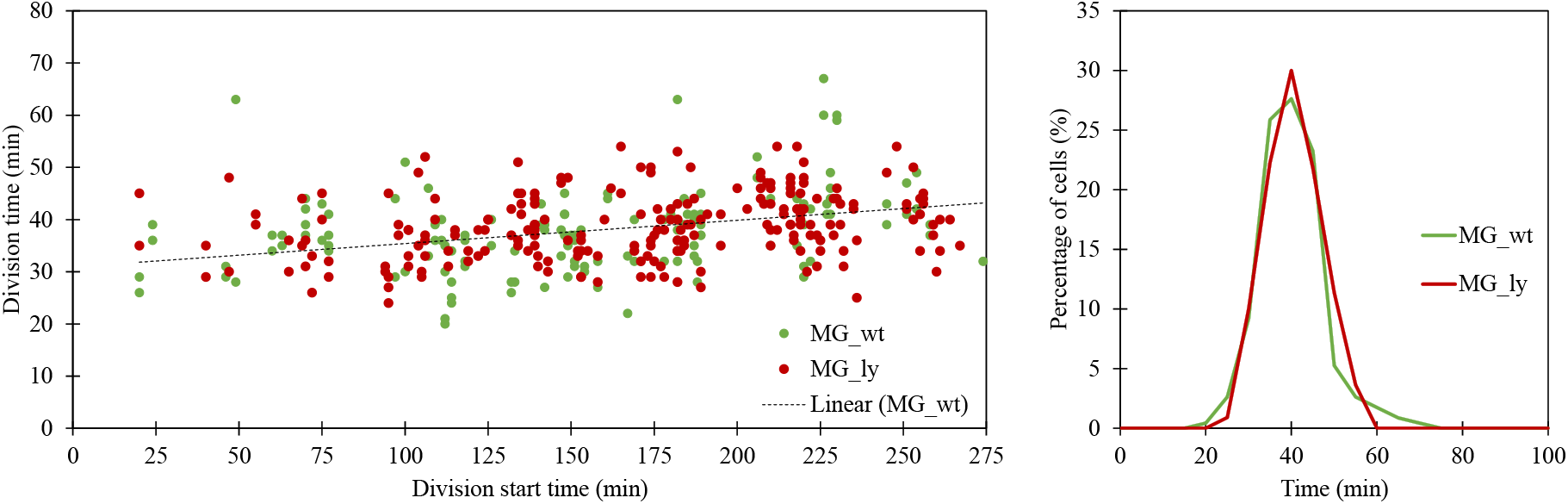
Division times of the wild-type and lysogenic cells. The data shows that the division times for the wild-type and lysogenic are nearly identical, *τ*_*d*_(wt) = 38.1± 7.4 mins and *τ*_*d*_(ly) = 38.7± 6.4 mins.

**Figure S4:**
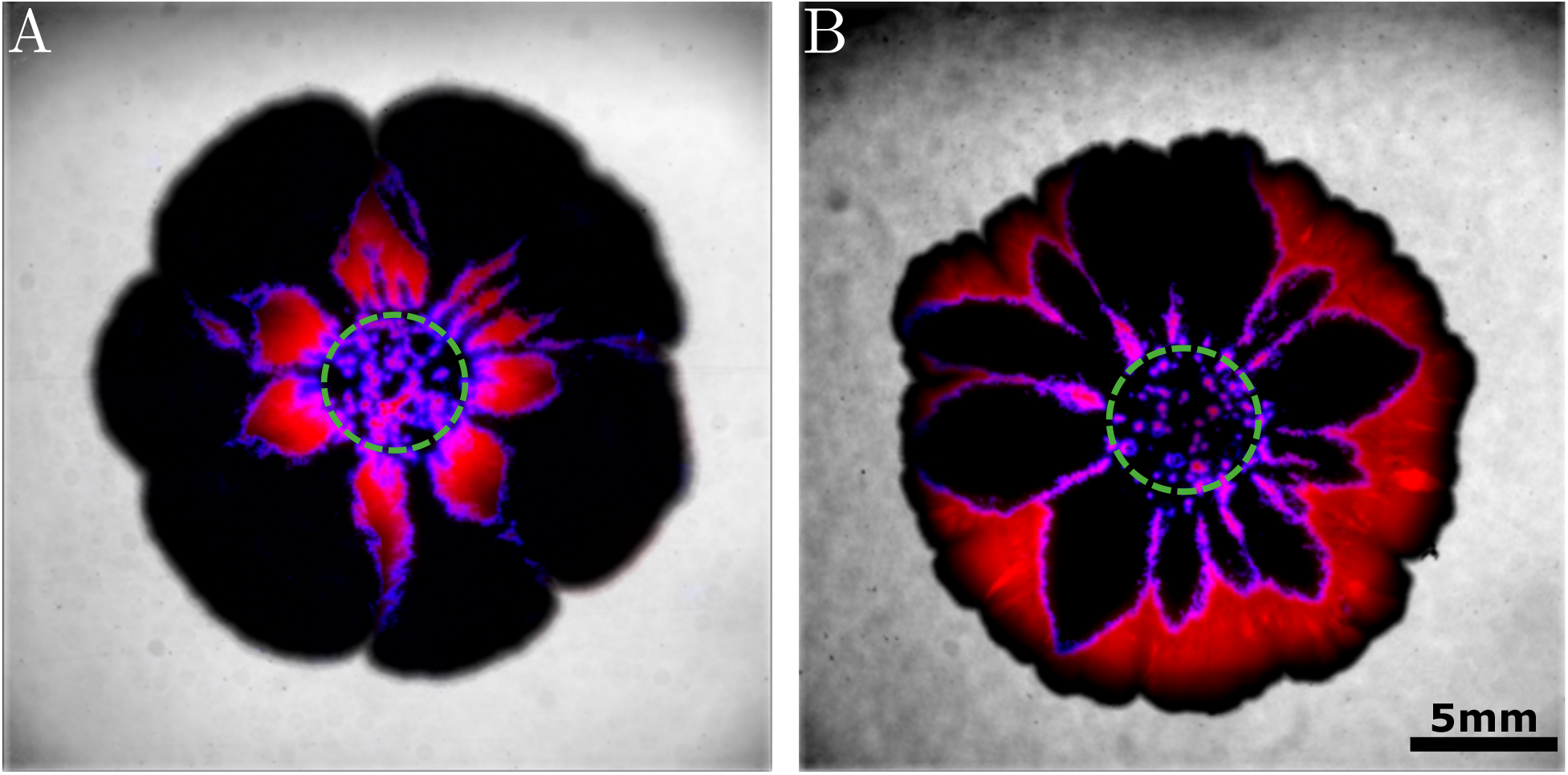
Diversion in outcome of colony front. **A**. In some cases we observe the uninfected cells overtake and encapsulate the lytic river, leading to extinction of the lysogenic front. **B**. In other cases the infection spreads along the growing front, hindering its growth. Allowing for the lysogenic sector to overtake.

**Figure S5:**
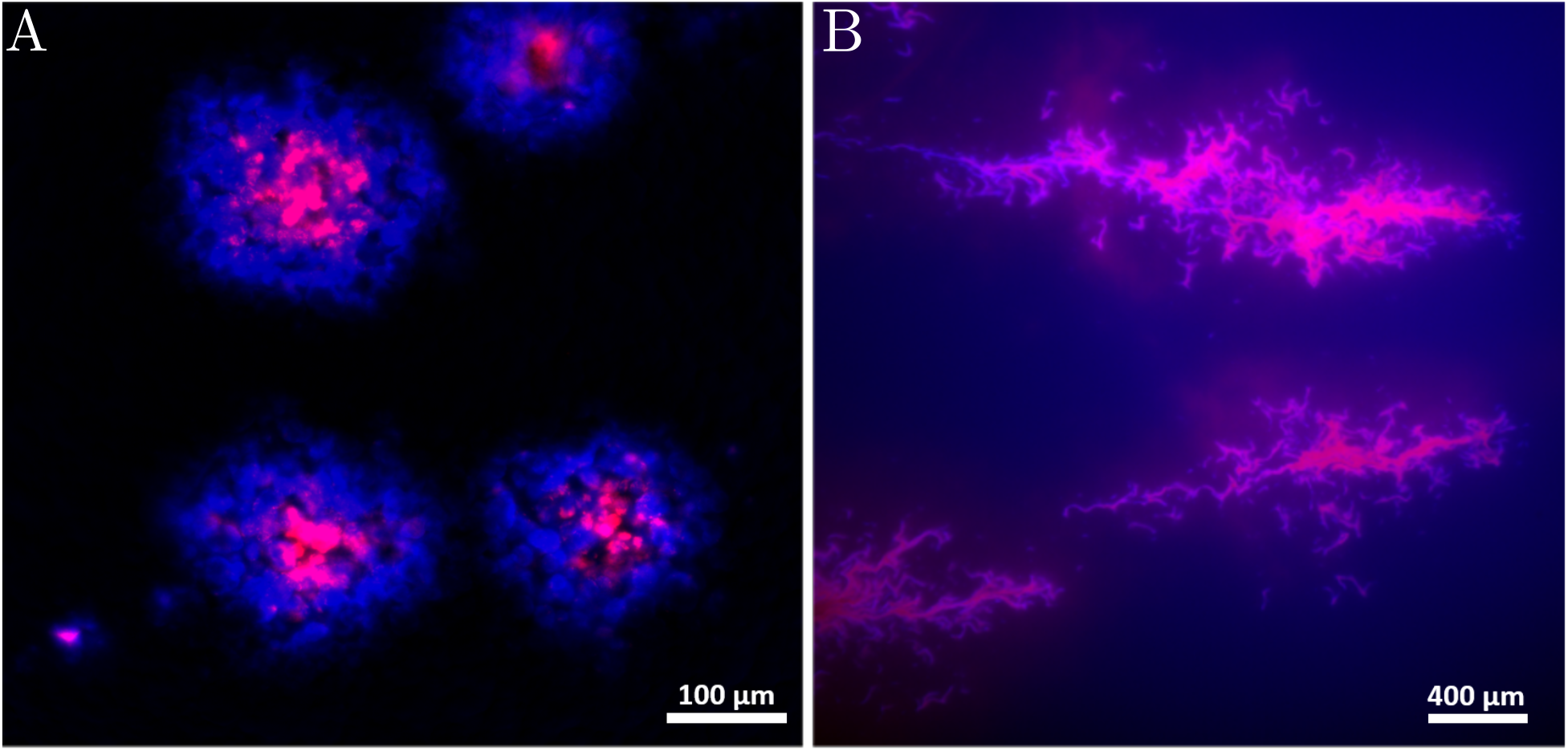
**A**. Regular plaques formed due to temperate phage infection on a lawn of bacteria under fluorescence microscope **B**. Lytic streams extinguished as fast growing uninfected cells flank the slow infected cells (Blue background is an artifact of channel bleed-through).

**Figure S6:**
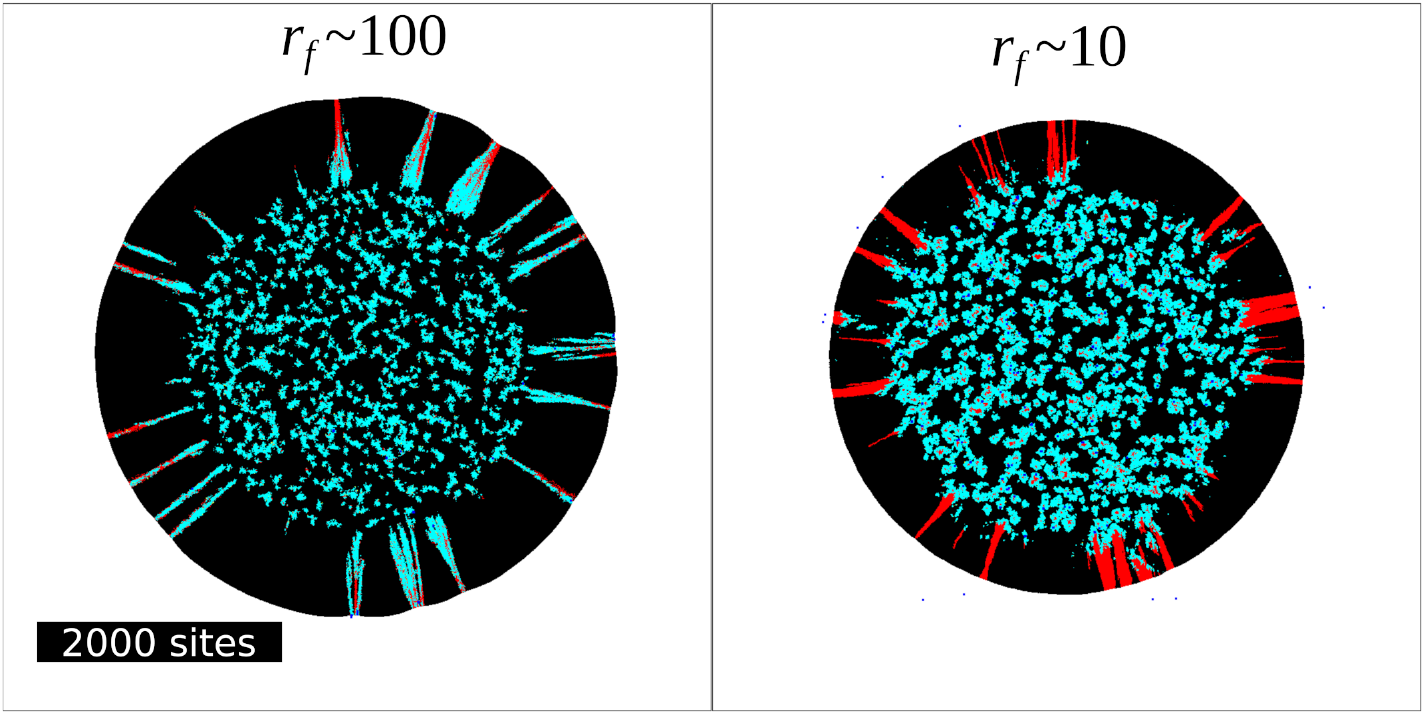
Effect of the advection by growth on the pattern. In both images, *p*_*diff*_ = 937.5, and initial *API* = 0.010. The radius for growth *r*_*f*_ is varied by 10-fold. Since this changes the colony expansion speed, similar size colonies are compared (the left panel is at time 32, the right panel is at time 192.) The phage diffusion parameter *p*_*diff*_ is set to a relatively small value to prevent the phages to cover the all the colony edge before colony grows significantly for small *r*_*f*_ case. Clearly, the larger *r*_*f*_, hence with more advection, the infection front have bigger chance of staying at the colony edge for longer length scale.

**Figure S7:**
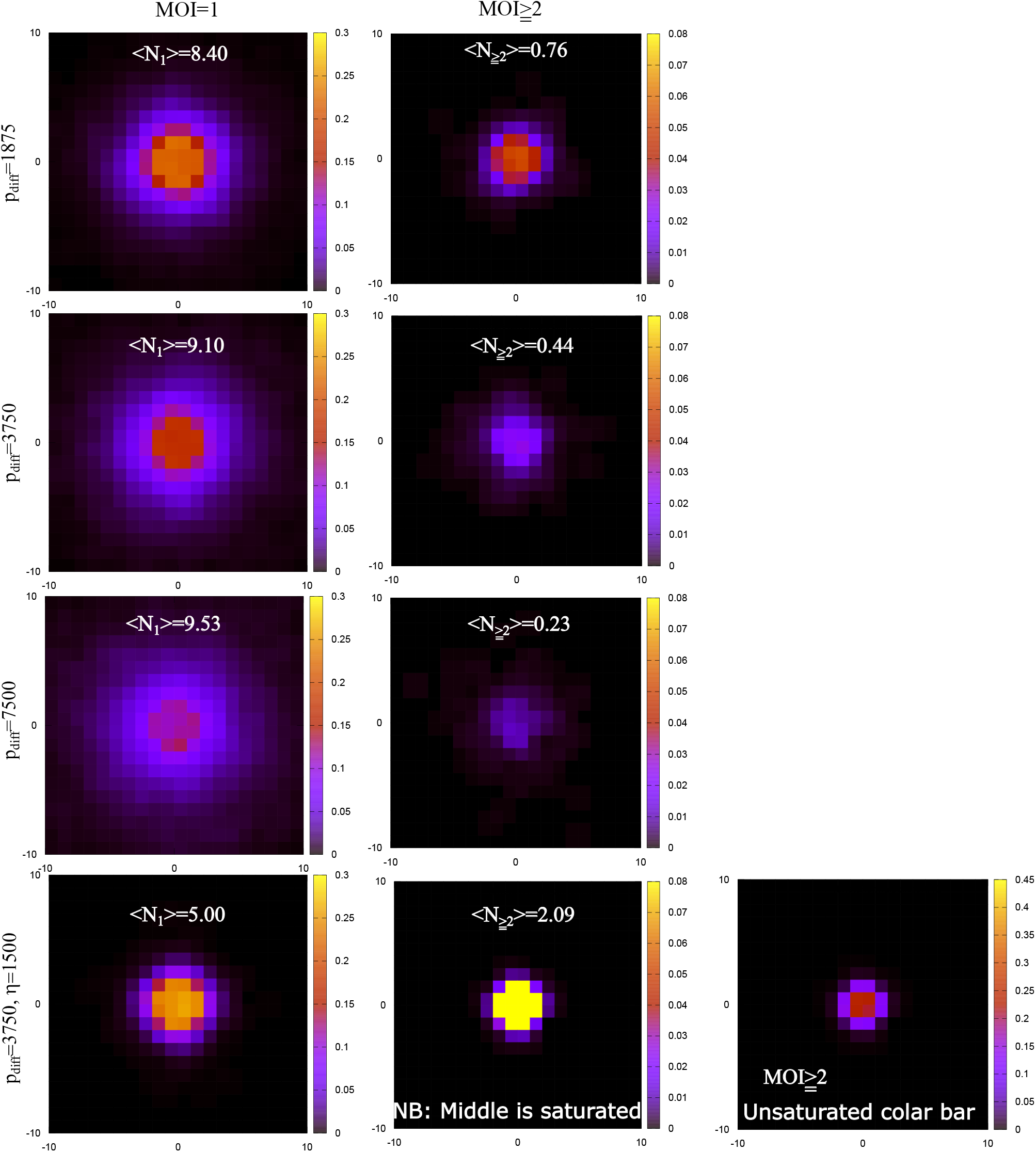
The multiplicity of infection after one burst in the middle of a dense colony. For the top 3 panels, *η* = 150, and *p*_*diff*_ is varied from 1875 to 7500. In the bottom panels, *η* = 1500 and *p*_*diff*_ = 3750. The left column shows the probability of being infected by 1 phage, and the right column shows the probability of being infected by 2 or more phages. *N*_1_ (*N*_*≥*2_) shows the sum of the probability over the system, resulting in the average number of *MOI* = 1 infections (*MOI ≥* 2 infections) in the system per burst.

